# Genome-wide histone modification patterns in *Kluyveromyces Lactis* reveal evolutionary adaptation of a heterochromatin-associated mark

**DOI:** 10.1101/039776

**Authors:** Angela Bean, Assaf Weiner, Amanda Hughes, Eyal Itskovits, Nir Friedman, Oliver J. Rando

## Abstract

The packaging of eukaryotic genomes into nucleosomes plays critical roles in all DNA-templated processes, and chromatin structure has been implicated as a key factor in the evolution of gene regulatory programs. While the functions of many histone modifications appear to be highly conserved throughout evolution, some well-studied modifications such as H3K9 and H3K27 methylation are not found in major model organisms such as *Saccharomyces cerevisiae*, while other modifications gain/lose regulatory functions during evolution. To study such a transition we focused on H3K9 methylation, a heterochromatin mark found in metazoans and in the fission yeast *S. pombe*, but which has been lost in the lineage leading to the model budding yeast *S. cerevisiae*. We show that this mark is present in the relatively understudied yeast *Kluyveromyces lactis*, a Hemiascomycete that diverged from *S. cerevisiae* prior to the whole-genome duplication event that played a key role in the evolution of a primarily fermentative lifestyle. We mapped genome-wide patterns of H3K9 methylation as well as several conserved modifications. We find that well-studied modifications such as H3K4me3, H3K36me3, and H3S10ph exhibit generally conserved localization patterns. Interestingly, we show H3K9 methylation in *K. lactis* primarily occurs over highly-transcribed regions, including both Pol2 and Pol3 transcription units. We identified the H3K9 methylase as the ortholog of *Set6*, whose function in *S. cerevisiae* is obscure. Functionally, we show that deletion of *KlSet6* does not affect highly H3K9me3-marked genes, providing another example of a major disconnect between histone mark localization and function. Together, these results shed light on surprising plasticity in the function of a widespread chromatin mark.

## INTRODUCTION

All genomic transactions in eukaryotes take place in the context of a chromatin template (KORNBERG and LORCH 1999). Chromatin plays key regulatory roles in control of transcription and other processes, and a great deal of cellular machinery is devoted to manipulation of nucleosome positioning (LI *et al.* 2007; JIANG and PUGH 2009; RADMAN-LIVAJA and RANDO 2010), subunit composition (HENIKOFF and AHMAD 2005), and covalent modification states (BERGER 2001; CAMPOS and REINBERG 2009; RANDO 2012).

Chromatin architecture is increasingly appreciated to play a major role in the evolution of gene regulatory programs. Most regulatory divergence between closely-related *S. cerevisiae* strains is associated with divergence in unlinked (*trans*) chromatin remodelers (BREM *et al.* 2002; LEE *et al.* 2006). Conversely, many transcriptional differences between *S. cerevisiae* and *S. paradoxus* (Last Common Ancestor (LCA) ~2 Million years ago (Mya)) are due to linked *cis* polymorphisms predicted to affect nucleosome occupancy (TIROSH *et al.* 2008; TIROSH *et al.* 2009). Over greater phylogenetic distances, nucleosomes have recently been mapped across 16 species of the *Ascomycota* phylogeny (TSANKOV *et al.* 2010; TSANKOV *et al.* 2011; XU *et al.* 2012), with evidence for numerous concerted shifts in chromatin structure and regulatory program. Most famously, changes in the regulation of mitochondrial ribosomal protein (mRP) genes between species that favor respiration vs. fermentation for energy production were correlated with changes in antinucleosomal PolyA-rich sequences – species that preferentially respire exhibit relatively long PolyA tracts at mRP promoters, which “program” a more open promoter structure at these genes (IHMELS *et al.* 2005; FIELD *et al.* 2009; TSANKOV *et al.* 2010).

While chromatin architecture at specific orthologous genes can differ between species, many global aspects of chromatin structure are conserved in all eukaryotes studied – the histone proteins are highly conserved at the amino acid level, for instance. Moreover, many covalent histone modifications are found in all organisms investigated, and exhibit conserved localization patterns with respect to underlying genomic features. For instance, trimethylation of histone H3 lysine 4 (H3K4me3) has been found in all eukaryotes investigated, and this mark is localized to the 5’ ends of transcribed genes in all species in which it has been mapped (RANDO and CHANG 2009; SHILATIFARD 2012). Although many histone modifications exhibit conserved localization patterns, their *functions* are not necessarily conserved. For example, while the “elongation” mark H3K36me3 is consistently localized over coding regions of genes, this mark serves primarily to suppress cryptic internal transcripts in budding yeast, whereas it is enriched over exons and plays a role in regulating splicing in metazoans (CARROZZA *et al.* 2005; KEOGH *et al.* 2005; LUCO *et al.* 2010).

In contrast to marks such as H3K4me3 and H3K36me3, other histone modifications are more phylogenetically restricted. Most notably, two major repressive marks are not found in the budding yeast *S. cerevisiae*. The polycomb-associated modification H3K27me3 is widespread in multicellular organisms, where it plays a key role in control of cell state inheritance (SCHUETTENGRUBER *et al.* 2007), but is absent in both *S. cerevisiae* and *S. pombe*. Similarly, methylation of H3K9, a modification central to heterochromatin formation and control of transposable elements (GREWAL and MOAZED 2003), is found in most metazoans and in *S. pombe*, but is absent in *S. cerevisiae*. Compared to nucleosome positioning (TSANKOV *et al.* 2010; TSANKOV *et al.* 2011; XU *et al.* 2012), histone modification patterns have been relatively understudied in comparative genomics studies.

We sought to gain a deeper understanding of the evolutionary dynamics associated with the loss of H3K9 methylation in *S. cerevisiae. S. cerevisiae* is a member of the *Saccharomycotina* yeast subphylum (Figure 1A), which includes species diverging before and after a key whole genome duplication event (KELLIS *et al.* 2004) (WGD) that marked a shift from using respiration for energy production (in pre-WGD species) to primarily using fermentation (in post-WGD species) (CONANT and WOLFE 2007). Several observations suggest that H3K9 methylation might be found in pre-WGD Hemiascomycetous species. First, RNA interference (RNAi), which plays a key role in H3K9 methylation in *S. pombe* (VOLPE *et al.* 2002), occurs in a subset of budding yeasts, with its loss in *S. cerevisiae* being linked to its incompatibility with the “killer” double-stranded RNA virus (DRINNENBERG *et al.* 2009; DRINNENBERG *et al.* 2011). It is currently unknown whether H3K9 methylation loss in *S. cerevisiae* occurred in concert with loss of RNAi, or whether these epigenetic systems were lost independently. Second, the H3K36 demethylase Rph1 in *S. cerevisiae* also exhibits vestigial H3K9 demethylation activity in vitro (KLOSE *et al.* 2007). As this demethylase has been linked to control of genes involved in carbohydrate metabolism (TSANKOV *et al.* 2010), we hypothesized that H3K9 methylation might play a role in regulation of these genes in pre-WGD species.

**Figure 1.**
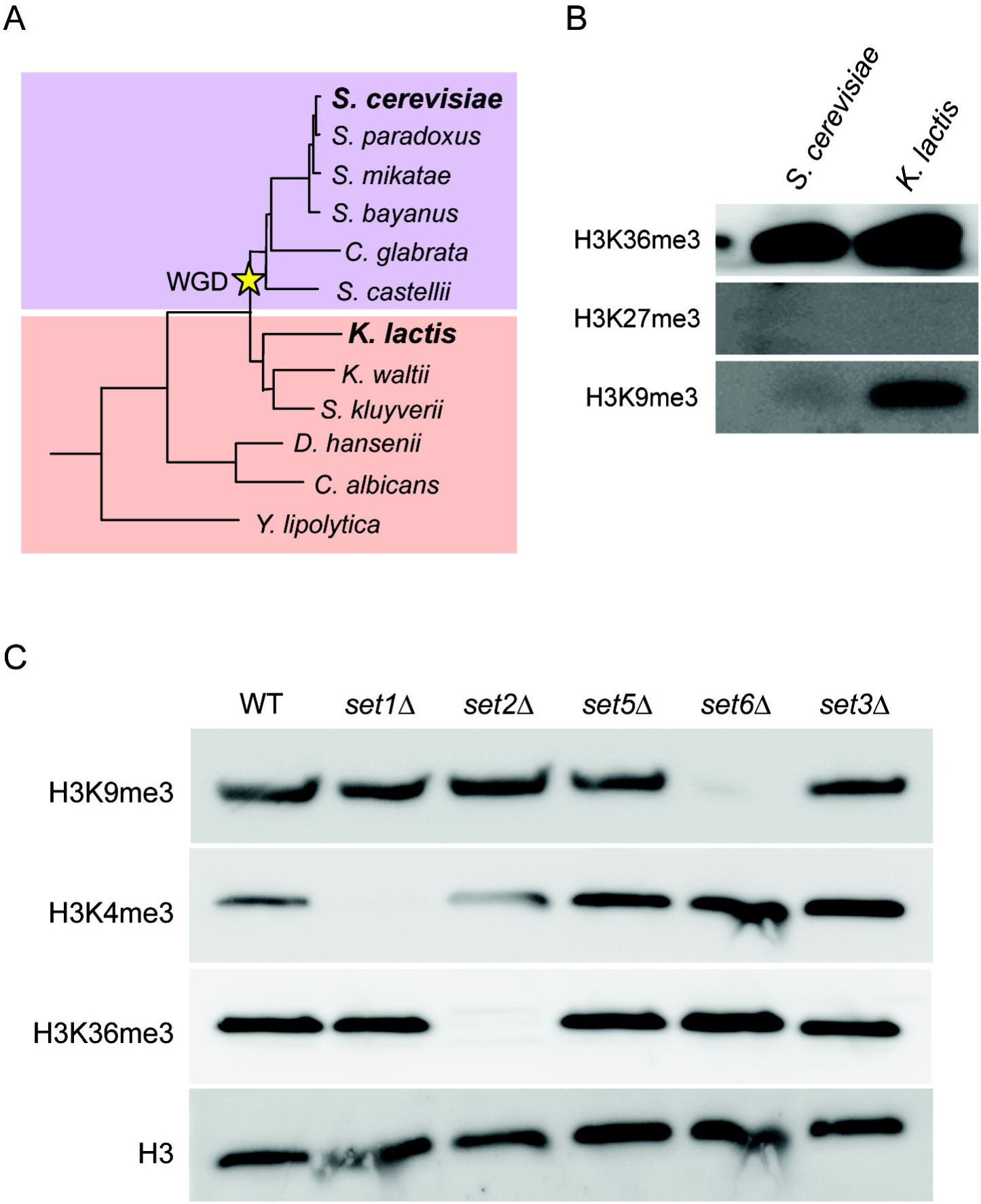
H3K9 methylation in *K. lactis*. **A**) Schematic of the Hemiascomycota phylogeny, adapted from Tsankov *et al* 2010. The key whole genome duplication event is marked with a star. The two species considered in this study are indicated in bold. **B**) Western blots against various histone modifications in *S. cerevisiae* and *K. lactis*, as indicated. Blots show the expected presence of H3K36me3 (and H3K4me3 – not shown) in both species, and the absence of H3K27me3 in both. Unexpectedly, H3K9 methylation is observed in *K. lactis*. **C**) Identification of the *K. lactis* H3K9 methyltransferase. 5 out of 6 SET domain-containing genes were successfully deleted, and Western blots were performed in the deletion strains as indicated.

Here, we focus on *Kluyveromyces lactis* (last common ancestor with *S. cerevisiae*: ~150 MYa), which diverged prior to the WGD. Surprisingly, we found that H3K9 methylation occurs in this organism, and identified the modifying enzyme as the ortholog of the SET domain protein Set6 of *S. cerevisiae*. We characterized the *K. lactis* epigenome in some detail, generating whole genome maps of transcriptional start sites, and of well-studied histone marks including H3K4me3, H3K36me3, H3S10ph, as well as H3K9me1, me2, and me3. To our surprise, H3K9 di-and tri-methylation was abundant over highly transcribed open reading frames, and over tRNA genes. Functionally, loss of H3K9 methylation did not preferentially affect H3K9me2/3-marked genes, consistent with the general disconnect between histone modification location and function seen for many other modifications. Together, these results reveal a great deal of conservation in chromatin structure and function across the Ascomycota phylogeny, but also reveal extensive plasticity in the role of H3K9 methylation over evolutionary time scales.

## RESULTS

### Abundant H3K9 methylation in *K. lactis*

To investigate whether *K. lactis* histones carry any phylogenetically-restricted histone marks such as H3K9 or H3K27 methylation, we carried out Western blots of nuclear extracts from *S. cerevisiae* and *K. lactis*. We confirmed the presence of several universal histone modifications including H3K4 and H3K36 methylation, as well as several histone acetylation states (Figure 1B and not shown). In contrast, neither yeast species carried detectable levels of the Polycomb-related histone modification H3K27me3. However, to our surprise, significant levels of the heterochromatin mark H3K9me3 were found in *K. lactis*, but not *S. cerevisiae* (Figure 1B). As histone methylation is typically catalyzed by SET domain proteins, we therefore attempted to delete all six genes encoding SET domain proteins in the *K. lactis* genome. We successfully deleted five of the six SET domain encoding genes in *K. lactis*, and characterized histone modifications in these mutants by Western blot. Four of the five mutants maintained high levels of H3K9me3, but the deletion of *KLLA0C09944g* eliminated all detectable H3K9 methylation (Figure 1C). The protein encoded by this gene is 45% identical to the *S. cerevisiae* Set6 protein and is considered its ortholog (WAPINSKI *et al.* 2007), so we have renamed this gene *KlSET6*. Curiously, while both Set6 orthologs bear reasonable matches to SET domain motifs III and IV, in neither protein do we find the conserved SET motifs I (GXG) or motif II (YXG) – associated with Adomet binding and catalytic activity, respectively (CHENG *et al.* 2005) – at the conserved spacing (Figure S1). A nearby YXL motif can be found in both Set6 orthologs however, and GXG and YXG sequence motifs can also be found elsewhere in the two Set6 proteins, albeit with spacing relative to the other SET domain motifs that would presumably necessitate significant structural perturbations to this domain (Figure S1). It is will be interesting in future studies to identify the structural basis underlying Set6 methylation activity, and to understand why the *S. cerevisiae* ortholog has lost K9 methylation activity, or whether H3K9 methylation occurs in *S. cerevisiae* under conditions that have not yet been investigated.

### Transcription start site mapping

We next sought to characterize the genome-wide distribution of histone modifications in *K. lactis*. As many aspects of chromatin structure correlate with transcript structure (JIANG and PUGH 2009; RADMAN-LIVAJA and RANDO 2010), we first mapped transcription start sites (TSSs) in this species to enable more precise alignments of chromatin maps. TSS mapping was carried out as previously described (NI *et al.* 2010) – briefly, polyadenylated RNA was isolated from actively growing *K. lactis* cultures, and uncapped transcripts were treated with phosphatase to prevent their capture during deep sequencing library construction. Transcripts bearing a 5’ cap were then cloned and deep sequenced, revealing the TSS landscape for this organism during exponential growth.

Alignment of TSS sequencing data with previously published nucleosome mapping data revealed a stereotypical relationship, with transcription on average initiating ~30 nt inside the upstream edge of the +1 nucleosome (Figure 2A). This confirms our prior speculation (TSANKOV *et al.* 2010; HUGHES *et al.* 2012) that transcription initiates relatively deeper into the +1 nucleosome in *K. lactis* than in *S. cerevisiae*, where the TSS is positioned ~12 nt inside the +1 nucleosome (JIANG and PUGH 2009). In addition to this apparent change in the architecture of the preinitiation complex, we also found that 5’ untranslated regions (the TSS-ATG distance) of mRNAs are generally longer in *K. lactis* than in *S. cerevisiae* (Figure 2B), potentially indicating a greater role for translational regulation in this organism. Thus, the increased distance from nucleosome-depleted region (NDR) to coding region ATG previously reported for *K. lactis* relative to *S. cerevisiae* (TSANKOV *et al.* 2010) reflects both altered transcript length as well as altered TSS location with respect to nucleosome positioning.

**Figure 2.**
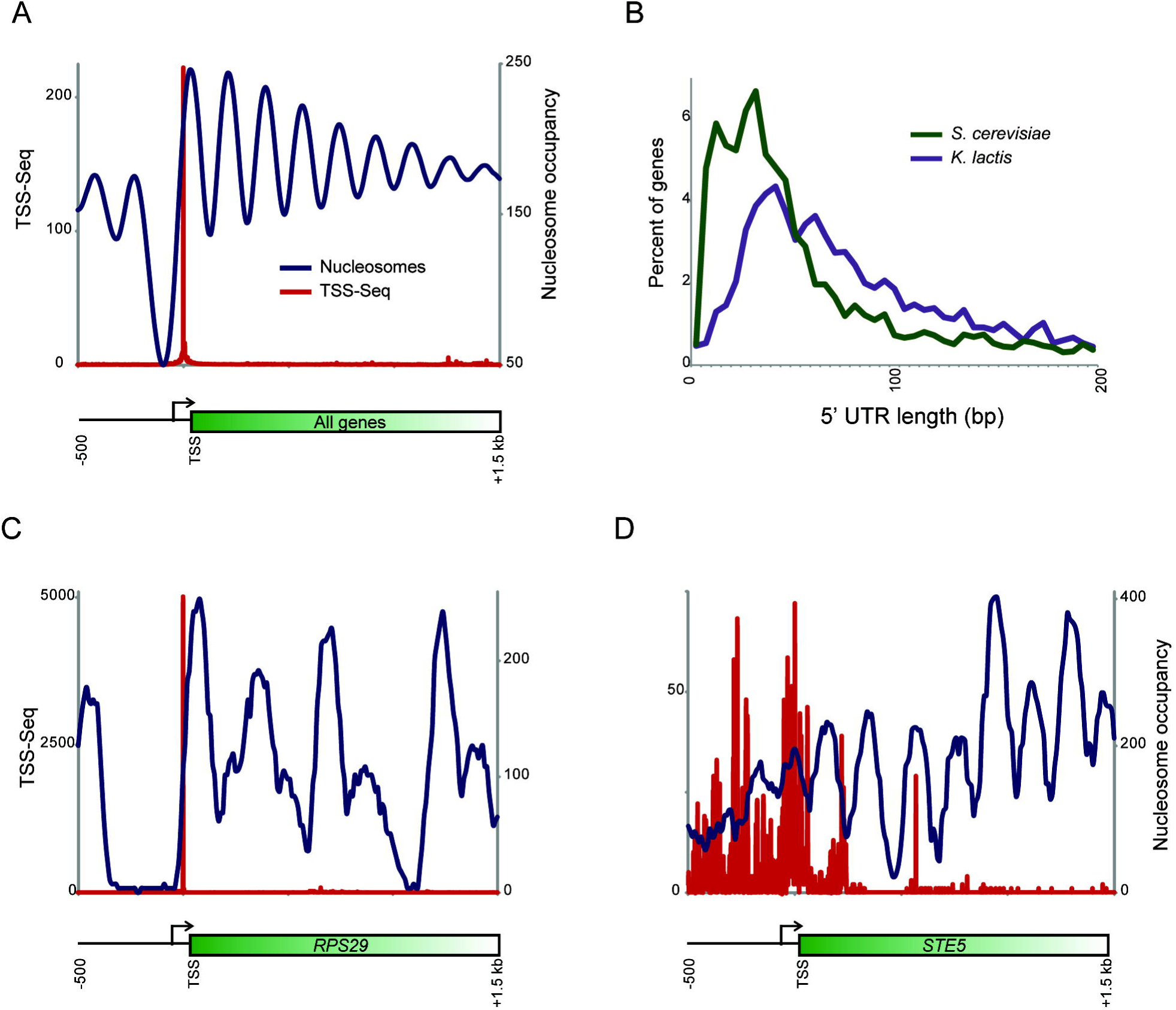
Transcription start site landscape in *K. lactis*. **A**) Transcription start sites (TSS) were characterized in actively growing *K. lactis* by deep sequencing. Here, data is shown averaged for all annotated open reading frames, aligned by the primary TSS identified in this dataset. TSS-Seq data is shown in red, and previous nucleosome mapping data (TSANKOV *et al.* 2010) is averaged and shown in blue. This representation reveals characteristic initiation of transcription inside the upstream border of the +1 nucleosome. **B**) Distribution of 5’ untranslated region (5’UTR) lengths for *K. lactis* and *S. cerevisiae*. 5’ UTR is defined as the distance from the primary TSS for a given gene to its start codon. **C-D**) Examples of genes with relatively unique (**C**) or dispersed (**D**) TSSs in *K. lactis*. TSS uniqueness was estimated here based on the fraction of all TSS-Seq reads contributed by the maximal peak, relative to all TSS-Seq data, for 1 kB surrounding each ORF ATG codon. Note that these genes are expressed at roughly similar overall levels when all sense TSS reads are summed.

Beyond the average relationship between TSS location and chromatin packaging, it is clear that the pattern of TSS usage differs between different classes of genes (RACH *et al.* 2011). We therefore defined for each TSS a “uniqueness” score based on the fraction of total TSS reads that were contributed by the major TSS peak. Sorting genes by this uniqueness score revealed that genes utilizing relatively few start sites were typically essential genes such as ribosomal protein genes (Figure 2C), whereas genes with broad groups of TSSs tended to be involved in processes such as pheromone response or protein folding (Figure 2D).

### Mapping of histone modifications – universal histone modifications

Having defined the landscape of TSS utilization for this species, we next turned to histone modification patterns. We first focused on the highly-conserved “initiation” mark H3K4me3 and the “elongation” mark H3K36me3, as well as the pericentric H3S10ph modification. We mapped these marks at nucleosome resolution – chromatin was digested to mononucleosomes using micrococcal nuclease, and chromatin immunoprecipitation using modification-specific antibodies was used to isolate nucleosomes bearing the relevant modifications (LIU *et al.* 2005). Enriched DNA was characterized by Illumina deep sequencing.

As shown in Figure 3A, H3K4me3 and H3K36me3 exhibit identical patterns to those seen in other organisms (LIU *et al.* 2005; POKHOLOK *et al.* 2005), with H3K4me3 enriched over 5’ ends of genes and H3K36me3 enriched over the middle and 3’ ends of genes. Both H3K4me3 and H3K36me3 levels are correlated with transcript abundance, whereas nucleosome occupancy is relatively uniform except at very high transcription rates (Figures 3B-D). H3S10ph localization was also consistent with prior data from *S. cerevisiae* (WEINER *et al.* 2012), as it was highly enriched over centromeres and over pericentric regions, and otherwise subtly anticorrelated with transcript abundance (Figure S2).

**Figure 3.**
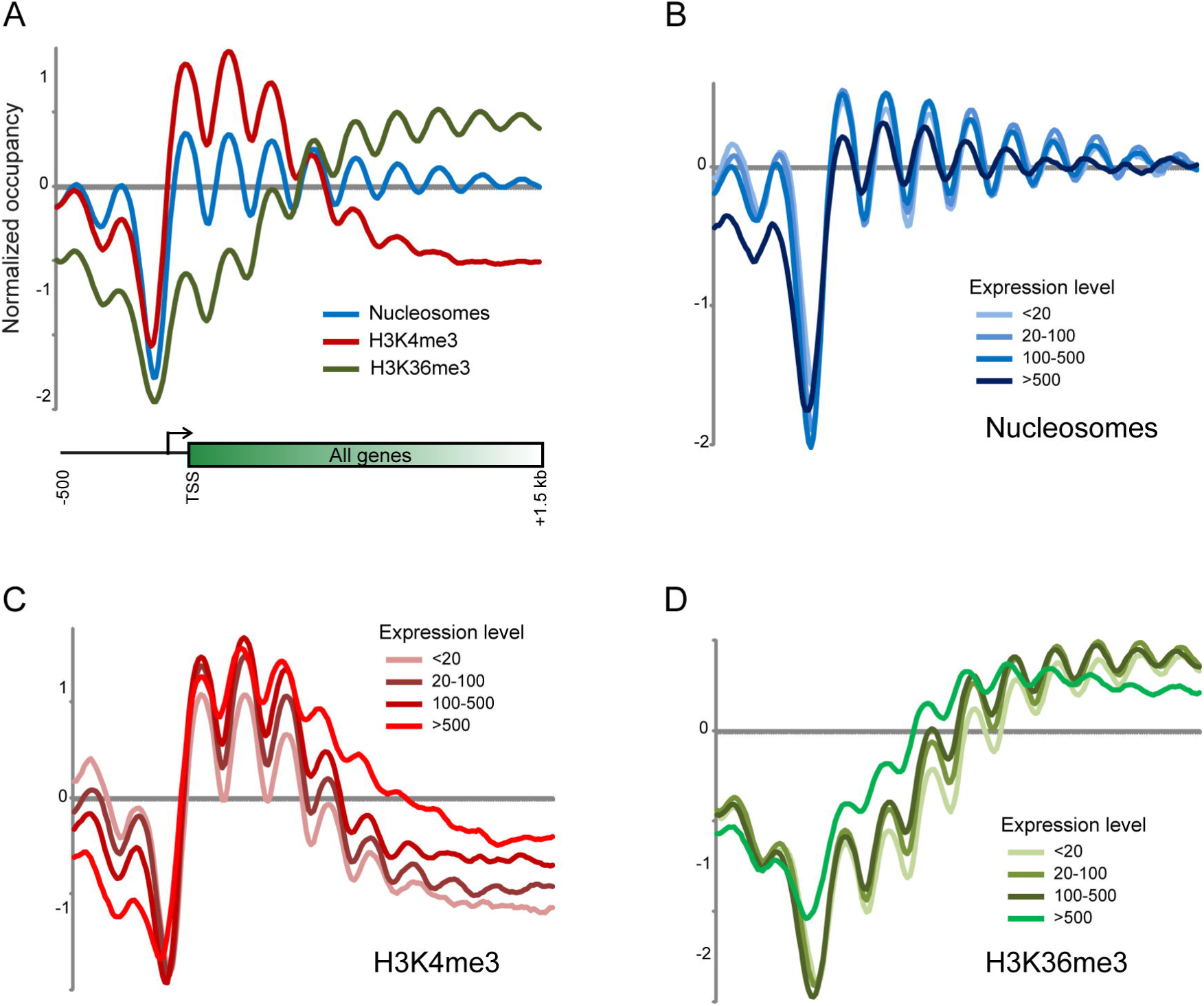
Conserved localization patterns for H3K4me3 and H3K36me3. **A**) Averaged data for all TSS-aligned genes for nucleosome mapping data and for H3K4me3 and H3K36me3 ChIP-Seq data, as indicated. **B-D**) Correlations between chromatin and transcription. For each mark, genes are broken into four groups based on mRNA abundance, as indicated. mRNA abundance is taken from Tsankov *et al*, 2010.

Thus, these three highly-conserved histone modifications are deposited by presumably conserved mechanisms in this understudied species. Nonetheless, we did find subtle differences between *S. cerevisiae* and *K. lactis* histone modification patterns. While H3K4me3 and other transcription-related marks are generally well-correlated with transcript abundance, there is nonetheless some variability in this relationship. By searching for genes that exhibit high levels of H3K4me3 or H3K36me3 *relative to their transcript abundance*, we found that cell cycle genes in *K. lactis*, but not in *S. cerevisiae*, exhibit higher histone methylation than expected given their mRNA abundance (not shown). This potentially reflects differences in cell cycle kinetics between these species – given that histone methylation over genes often persists after transcriptional shutdown until the following S phase (RADMAN-LIVAJA *et al.* 2010), a longer G1 phase would be expected to result in relatively longer perdurance of H3K4me3 for transiently transcribed genes.

These data provide a resource for future investigations into the chromatin structure of this understudied organism, and underscore the generally conserved behavior of many histone modifications.

### H3K9 methylation is associated with highly-transcribed loci in *K. lactis*

We next turned to genome-wide localization studies to characterize the distribution of H3K9 methylation across the *K. lactis* genome. Nucleosome resolution ChIP-Seq was carried out using antibodies against H3K9me1, me2, and me3. Unexpectedly, we found no particular enrichment of H3K9 methylation over classical heterochromatic regions such as transposable elements or over subtelomeric genes (not shown), instead finding surprisingly abundant H3K9me2 and me3 over protein-coding genes. Over open reading frames, H3K9 methylation was broadly localized over the middle and 3’ ends of genes (Figure 4A). H3K9me1 was generally enriched towards the 3’ ends of coding regions, while K9me2 and me3 were more abundant closer to the 5’ ends of genes. To our surprise, H3K9 tri-and di-methylation was generally more abundant over highly-transcribed genes – we found the highest levels of H3K9me3 and me2 over ribosomal protein genes, and on average H3K9me2 and me3 levels were correlated with transcript abundance (Figures 4B-D, Figure S3). More surprisingly, we also found peaks of H3K9me2 and me3 over tRNA genes – this is shown in aggregate for all tRNAs in Figure 5A, and is shown for all individual tRNA genes in Figure 5B. Taken together, our results reveal surprisingly divergent behavior of H3K9 methylation in *K. lactis*, as this classical mark of heterochromatin appears to have completely lost this aspect of its genomic localization.

**Figure 4.**
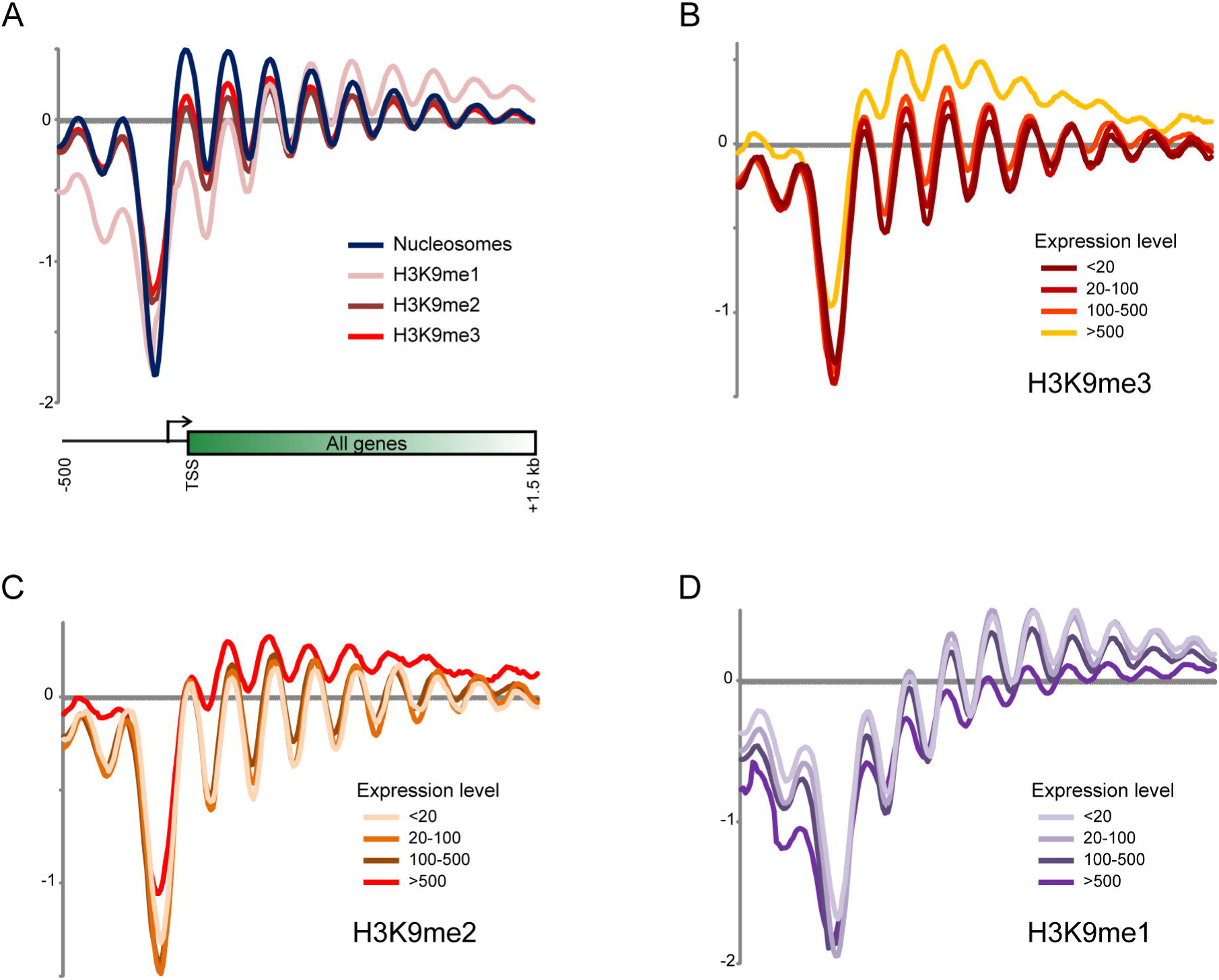
Localization patterns of H3K9 methylation states. **A**) Averaged data for all TSS-aligned genes for nucleosome mapping data and for the three H3K9 methylation states. **B-D**) Correlations between chromatin and transcription. As in **Figures 3B-D**, but for H3K9 methylation marks.

**Figure 5.**
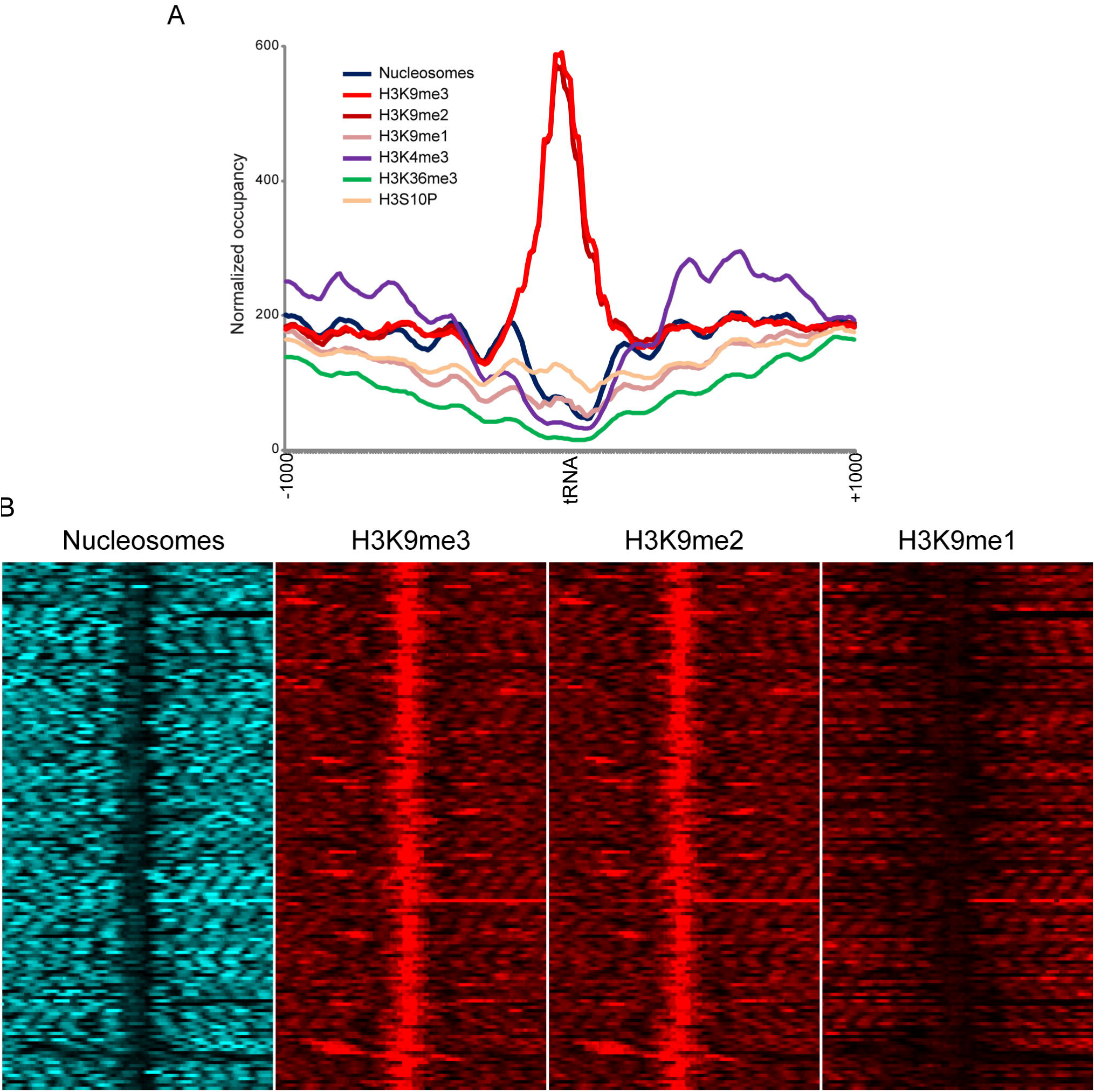
H3K9 methylation over tRNA genes. **A**) Data for all histone marks in this study are averaged across all tRNA genes along with 1 kb of flanking sequence. High H3K9me2 and me3 signal is clearly not an artifact of MNase sequence specificity, or of nonspecific antibody binding to genomic loci with low average protein content. **B**) Individual tRNA genes are shown in a heatmap for the indicated datasets, showing H3K9 methylation across all tRNA genes. Note that although bulk nucleosomes are depleted over tRNA genes, in all cases there is a low occupancy nucleosome observed which we infer based on the H3K9me3 ChIP-Seq to be H3K9 methylated at very high levels.

### Functional consequences of H3K9 methylation

The localization of H3K9 methylation over highly-transcribed genomic loci was quite unexpected given that this modification is a mark of heterochromatin in most species (GREWAL and MOAZED 2003). Given the widespread disconnect between histone modification localization and function – histone marks generally occur primarily at genomic loci that are unaffected by loss of the mark (RANDO 2012) – we sought to understand the regulatory role for H3K9 methylation in *K. lactis* transcription. First, to investigate whether *KlSET6* deletion grossly altered rDNA or tRNA transcription, we assessed the size distribution of total RNA isolated from wild-type and from *Klset6Δ K. lactis* strains. Overall, size distributions of the total RNA profile were nearly identical between wild-type and *Klset6Δ* strains (Figure S4), indicating no global effects of this mutant on rDNA or tRNA expression. Moreover, doubling time of the mutant strain was similar to that of the wild-type, indicating no dramatic consequences of loss of H3K9 methylation on strain growth.

RNA-Seq of polyadenylated transcripts was then used to more specifically characterize the effects of H3K9 methylation on mRNA abundance. As is often seen in chromatin regulatory mutants (LENSTRA *et al.* 2011; RANDO 2012), the global transcriptome was relatively insensitive to loss of all H3K9 methylation (Figure 6A). Most relevant in this context, we note that highly-expressed mRNAs in wild-type *K. lactis* remain highly-expressed in *Klset6Δ*. Consistent with this observation, genes associated with high levels of H3K9me2 or me3 in wild type cells were generally transcriptionally unaffected in the *Klset6Δ* – genes mis-regulated in this mutant in fact had relatively *low* levels of H3K9me2 and me3, and average levels of H3K9me1, in wild-type *K. lactis* (Figure 6B and not shown). As is typical for genome-wide studies, it is worth noting that this result follows genome-wide normalization, and thus reflects effects of gene expression *relative* to the bulk of the transcriptome, but as shown in Figure S4 overall mRNA abundance was grossly unaltered in the absence of H3K9 methylation.

**Figure 6.**
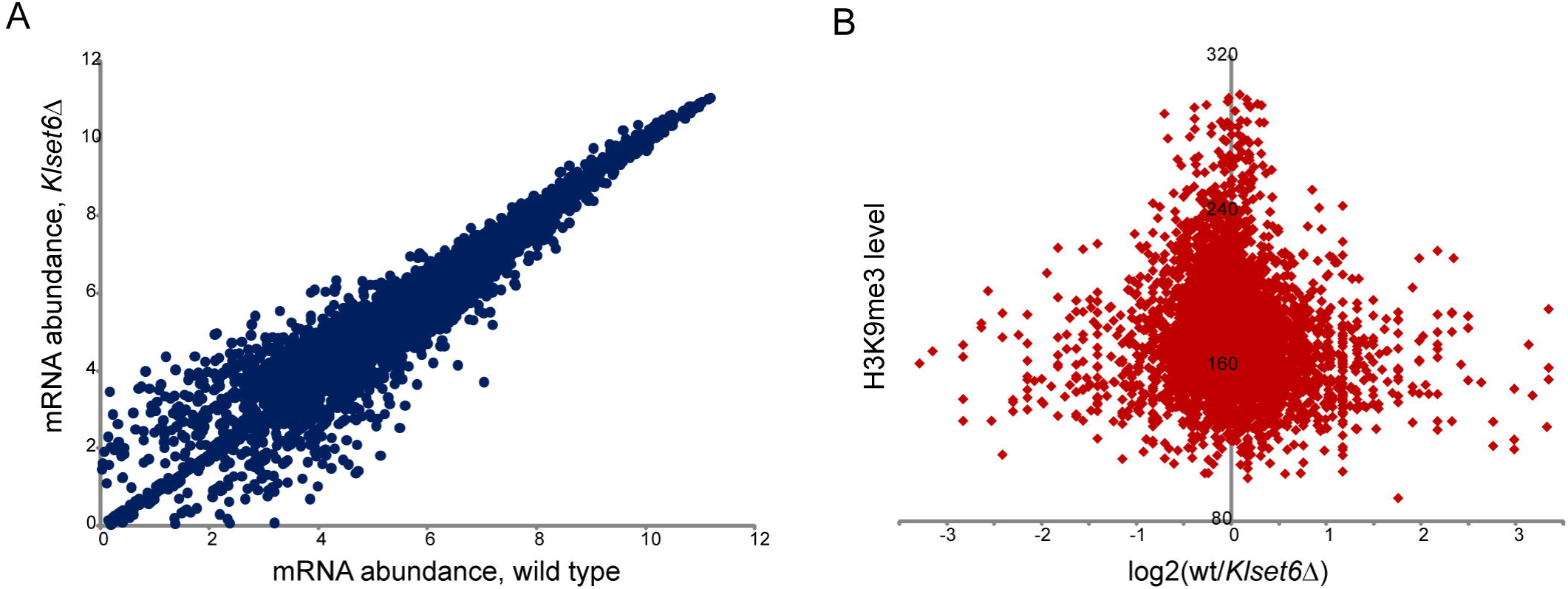
Function of H3K9 methylation. **A**) Scatterplot of mRNA abundance from wild-type and *Klset6*Δ strains, as indicated. Axes show normalized read count as a log2 value. **B**) Comparison between H3K9me3 levels over coding regions and change in mRNA abundance in *Klset6*Δ yeast. Genes marked with high levels of H3K9me3 are not preferentially affected by loss of global H3K9me3. Similar results hold, but are not shown, for H3K9me2 and H3K9me1.

While misregulated genes were not generally associated with high levels of H3K9 methylation, they did show significant enrichments for particular functional gene categories. Specifically, genes repressed in the deletion mutant included a large number of genes predicted to play roles in splicing (p < .001 for genes downregulated at least 2-fold). In contrast, upregulated genes did not exhibit significant GO enrichments. Taken together, these results reveal that the function of H3K9 methylation in gene regulation is disconnected from its genomic localization pattern, consistent with results found for the majority of other histone modifications.

## DISCUSSION

Here, we explored the genomic histone modification landscape of the understudied yeast species *Kluyveromyces lactis*. For the majority of histone modifications analyzed, we found that the genomic landscape of the modification was broadly concordant with localization patterns previously reported in various other species. Specifically, H3K4me3 and H3K36me3 were associated with regions of RNA Pol2 initiation and elongation, respectively, while H3S10ph was primarily localized to pericentromeric regions (presumably largely during M phase) and also was subtly anti-correlated with transcription rate over coding regions. These evolutionarily conserved *localization* patterns do not necessarily imply conservation of *function* – as noted in the introduction, H3K36 methylation impacts splicing in metazoans, versus cryptic promoter repression in budding yeast – but more likely they reflect conserved mechanisms of mark deposition. In other words, H3K4me3 appears to be universally deposited by a methyltransferase associated with the initiation form of RNA Pol2, and thus serves as a universal mark for promoters and genic 5’ ends. In contrast, given the modular nature by which histone modifications recruit or activate effector complexes through specific modification-binding domains, the functional role for this 5’ gene mark can readily change over evolutionary time. While this view significantly oversimplifies the true biology of a given histone modification – gene duplication and divergence often leads to a number of histone modifying enzymes specific for a given residue being found in a species – it serves to reinforce the idea of a core evolutionarily-conserved deposition machinery that may be further elaborated upon.

### Novel roles for H3K9 methylation

The most interesting facet of this study is the discovery of H3K9 methylation in a budding yeast. H3K9 methylation is an extensively-studied histone modification that is generally associated with epigenetic gene repression, primarily of selfish genetic elements such as transposons. Among fungi, H3K9 methylation has been extensively studied in the fission yeast *S. pombe*, where it plays key roles in silencing of the silent mating loci and of pericentric heterochromatin and in “facultative heterochromatin” (as found for example at genes encoding meiosis-specific proteins (ZOFALL *et al.* 2012)). A multitude of mechanisms targeting H3K9 methylation have been described, including targeting by small interfering RNAs (siRNAs), targeting to meiotic transcripts by an RNA-binding protein specific for a particular hexanucleotide motif, targeting via sequence-specific DNA-binding domains, and targeting by tethering of HP1 to nascent transcripts. While H3K9 methylation is typically a mark of heterochromatin, it also can be found over euchromatic genes where its functions are far more poorly understood. In flies, HP1 and H3K9 methylation are both found associated with active genes located within heterochromatin (as for example on the heterochromatic “dot” chromosome 4), and HP1 is also associated with exon-dense active genes in euchromatin where it may play a role in stabilization of a subset of mRNAs (DE WIT *et al.* 2007; JOHANSSON *et al.* 2007; PIACENTINI *et al.* 2009).

However, H3K9 methylation is not universally found in eukaryotes – most notably, it has apparently been lost in *S. cerevisiae*, where analogous silencing is performed by the Sir H4K16 deacetylase complex (RUSCHE *et al.* 2003). To our knowledge, H3K9 methylation has not been previously observed in any Hemiascomycete yeast. Here, we show for the first time the presence of H3K9 methylation in a Hemiascomycete, and show loss of detectable H3K9 methylation in mutants lacking the ortholog of *S. cerevisiae SET6*. Although we have not carried out in vitro histone methylation assays with Set6p and therefore cannot state unequivocably that Set6 is the histone methylase in this system, the complete loss of H3K9me3 in this mutant, coupled with the fact that SET domain enzymes carry out most histone lysine methylation, make this clearly the simplest interpretation of our results. It is unclear at present whether Set6 in *S. cerevisiae* might be capable of H3K9 methylation under unusual growth conditions, as both species’ Set6 orthologs lack conserved spacing of certain key sequence motifs, meaning that at present the relevant sequence changes between species cannot be interrogated by sequence analysis.

Given the generally conserved role of H3K9 methylation in heterochromatin, our finding of abundant H3K9me2 and me3 over highly-transcribed genes – both mRNAs and tRNAs – was unanticipated, although as noted above H3K9 methylation can also be found associated with active genes in some species. In *K. lactis*, we find that genomic loci marked with high levels of H3K9me2 or me3 exhibit two major features – they are highly-transcribed, and the majority of these loci produce some noncoding RNA. tRNA genes are of course entirely noncoding, while the most highly-expressed mRNAs in *K. lactis* under our growth conditions are the ribosomal protein genes, which in this species as in *S. cerevisiae* are among the rare genes that carry introns. How is KlSet6-dependent H3K9 methylation targeted? *K. lactis* appears to lack the Dcr1-type RNaseIII homolog required for siRNA production (DRINNENBERG *et al.* 2011), suggesting that H3K9 methylation is likely to be targeted in this species by some other aspects of noncoding transcripts, or potentially by a novel mechanism such as binding of the KlSet6 protein to elongating RNA polymerases.

Finally, evolutionary divergence in the function of this modification will be interesting to consider in this species. In this regard it is interesting that Dicer-mediated siRNA production has been shown to occur in only a subset of Ascomycota (such as *S. pombe* and *S. castelii*, but not in *S. cerevisiae*), and the loss of this pathway has been linked to its effects on the function of the double-stranded RNA killer virus (DRINNENBERG *et al.* 2011). In *K. lactis*, loss of H3K9 methylation affects mRNA abundance of genes encoding components of the spliceosome. Given the functional role for another RNase III, Rnt1, in processing of noncoding RNAs such as those involved in the spliceosome, it is tempting to speculate that H3K9 methylation in *K. lactis* affects noncoding RNA processing (LEE *et al.* 2013), with spliceosomal mRNA abundance changing as a secondary consequence of altered ncRNA processing. In any case, the pressures resulting in the altered localization and function of H3K9 methylation in this species may provide an interesting test case for evolutionary lability in the function of the epigenome.

## MATERIALS AND METHODS

### Growth Conditions and strains

S. *cerevisiae* and K. *lactis* strains were grown in medium containing: Yeast extract (1.5%), Peptone (1%), Dextrose (2%), SC Amino Acid mix (Sunrise Science) 2 g/liter, Adenine 100 mg/L, Tryptophan 100 mg/L, and Uracil 100 mg/L. This medium was described previously in Tsankov et al. (2010).

K. *lactis* deletion strains were constructed using homologous recombination that replaced the gene of interest with a kanamycin resistant gene. Briefly, constructs were made with approximately 200 bp of homologous sequence around the gene of interest on each side of the kanamycin resistant gene. 1 µg of the linear construct was transformed into yeast and colonies were selected for kanamycin resistance. Deletions were confirmed using PCR.

Unfortunately, during the preparation of this manuscript, our stock of the *Klset6Δ* strain was accidentally lost. As the personnel associated with this project are now pursuing other efforts, this precludes us from carrying out one key control – mapping of H3K9 methylation patterns in *K. lactis* strains lacking this mark, to ensure that the antibodies used are indeed specific for the modifications in question. For this reason we are posting this manuscript to a preprint server in the hopes that the analyses here will assist researchers interested in this topic.

### Western Blotting

Cultures of wild type K. *lactis* and the deletion strains were grown to mid-log phase. 10 mLs of the culture was spun down, flash-frozen in liquid nitrogen, and then resuspended in lysis buffer containing 50mM Tris pH 6.8, 10% glycerol, 144mM Beta-mercaptoethanol, 2mM EDTA, 2% SDS and protease inhibitors. Then 0.5mM beads were added and subjected to 3 rounds of beating with 20s on and 20s off. The supernatant was collected in a new tube and used for subsequent western blotting. Antibodies used for western blotting: Anti-histone H3 (tri methyl K27) antibody (Millipore, 07-449) Anti-histone H3 (tri methyl K9) antibody (abcam, ab8898) Anti-histone H3 (tri methyl K36) antibody (abcam, ab9050-100)

### TSS Mapping

Trizol-extracted RNA was isolated from mid-log cultures and enriched for mRNA on polyT magnetic beads. The polyadenylated RNA was then treated with bacterial alkaline phosphatase (TAKARA) to remove all phosphates. The mRNA cap was then hydrolyzed to a phosphate by treating the RNA with Tobacco Acic Pyrophosphatase (Epicentre). This allowed for the subsequent ligation (T4 RNA ligase) of an oligo containing an MmeI site selectively to the 5’ end of RNA that had been previously capped. This RNA was reverse transcribed (SuperScript III Reverse Transcriptase, Invitrogen) with a tailed random primer. The cDNA was amplified with a low-cycle PCR (Phusion, NEB) using biotin-linked primers that matched the sequences added in the ligation step. An MmeI digestion then allowed for a cut site to be made 20bp downstream from where the 5’ cap had been. This DNA was then isolated with streptavidin beads and ligated to a modified illumina adaptor. After elution from the beads, a final PCR amplified the TSS sequences which were then cloned and deep sequenced.

### ChIP-Seq

Antibodies used:

Anti-histone H3 (mono methyl K9) antibody (abcam, ab9045)

Anti-histone H3 (di methyl K9) antibody (abcam, ab1220)

Anti-histone H3 (tri methyl K9) antibody (abcam, ab8898)

Anti-histone H3 (tri methyl K36) antibody (abcam, ab9050-100)

Anti-histone H3 (tri methyl K4) antibody (abcam, ab1012)

Anti-histone H3 (Serine 10 phosphorylation) antibody (Millipore, 04-817)

Briefly, 500mL of cells were grown to mid-log phase at 30°C, 220rpm. Cells were then fixed for 30 minutes at 30°C with a final concentration of 1% formaldehyde then spheroplasted with 10 mg of zymolyase for 55 minutes at 30°C. Spheroplasts were digested with micrococcal nuclease (MNase), the optimal concentration was determined through a prior titration, for 20 minutes. The digested product was then pre-cleared with a protein-A bead slurry prior to immuno-precipitation with protein A beads along with the indicated antibody overnight at 4°C with rotation. ChIP DNA was then isolated, blunt ended (Epicentre End-it repair kit), A-tailed (Epicentre–Exo Klenow), ligated to illumina adaptors, and amplified through PCR.

Note that H3K9me2 and me3 profiles were extremely well-correlated (Figure 4). This suggests that these antibodies are potentially poorly specific for me2 vs. me3, or that these marks broadly co-occur. However, none of the conclusions of this study depend on this distinction, and all signal for the H3K9me3 antibody is lost in *Klset6*Δ strains (Figure 1C). As a result we simply note this potential antibody concern here.

### Strand Specific RNA-Seq

Total RNA was isolated from mid-log cultures of K. *lactis* and the deletion strains using Trizol followed by a Phenol-Chloroform extraction and ethanol precipitation. Polyadenylated RNA was enriched using PolyT magnetic beads and first-strand cDNA synthesis was carried out using random hexamers and SuperScript III (Invitrogen). After a phenol-chloroform extraction followed by an ethanol precipitation a second strand cDNA synthesis reaction was carried out with E. *coli* DNA polymerase, E. *coli* DNA ligase and E. *coli* RNase H. The cDNA was purified with Zymo clean and concentrator kit. Fragmentation of the cDNA was done using the Fragmentase enzyme (NEB, #M0348L) and the fragments of interest were purified using Ampure XP beads (Beckman, #A63881). DNA was then isolated, blunt ended (Epicentre End-it repair kit), A-tailed (Epicentre–Exo Klenow), ligated to illumina adaptors, amplified through PCR then purified using Ampure XP beads.

## ACKNOWLEDGEMENTS

We thank H. Chen for assistance with TSS mapping, B. Carone for assistance with data analysis, and P. Kaufman and members of the Rando lab for critical discussions and comments on the manuscript.

---

**Figure S1.**
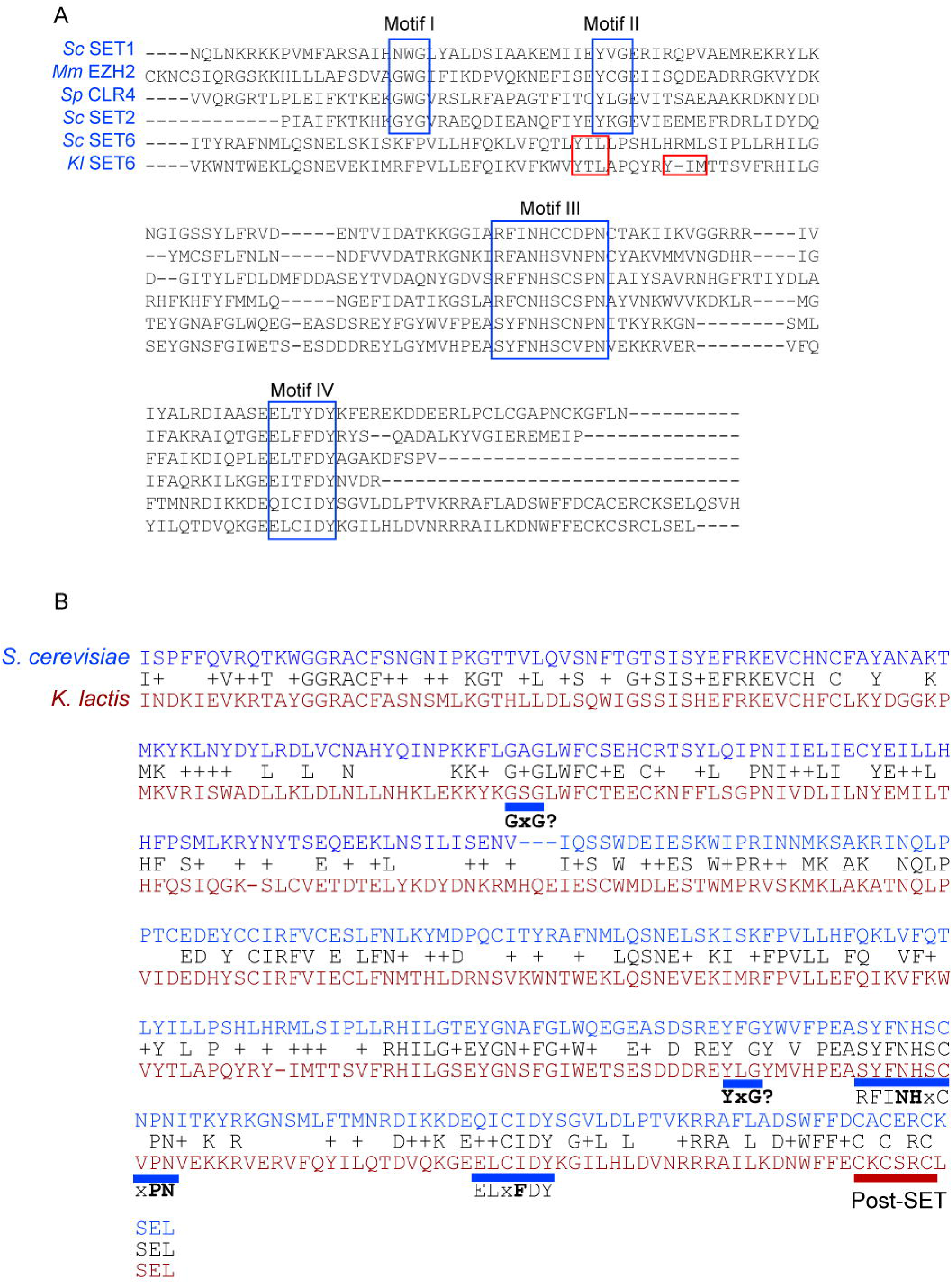
Set6 alignments. **A**) Multiple protein alignment using the SET domains from *S. cerevisiae* Set1, Set2, and Set6, *S. pombe* Clr4, *M. musculus* Ezh2, and *K. lactis* Set6. For the top four proteins, conserved Set domain motifs I, II, III, and IV are boxed in blue. As Motifs I and II are not found at the conserved spacing in either Set6 protein, we also indicate potential catalytic tyrosines in red boxes for Set6 proteins. **B**) Pairwise alignment of the two Set6 proteins studied here. Alternative motifs I and II are indicated. A cysteine-rich post-SET domain is also shown with a red underscore.

**Figure S2.**
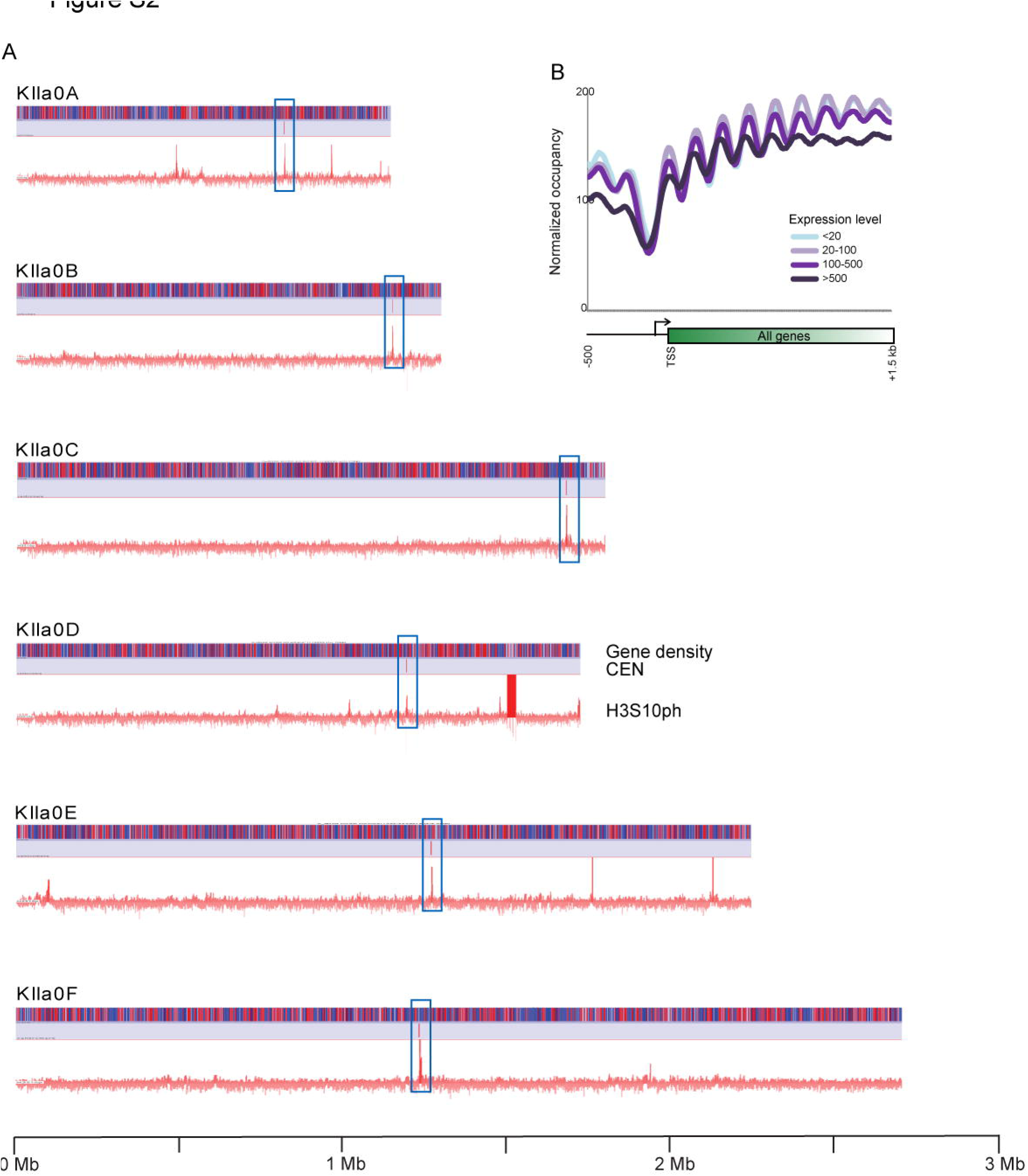
H3S10 phosphorylation in *K. lactis*. **A**) Chromosome views of gene density (top bar), centromere location (middle bar), and H3S10ph ChIP-Seq signal (bottom bar). Every centromere (highlighted with a blue box) is associated with a large peak of H3S10ph, consistent with results obtained in *S. cerevisiae* and other organisms. A small number of additional peaks of this size are observed, which occur over an interesting group of genes – predicted to be involved in ergosterol biosynthesis, in fatty acid beta oxidation, in the pachytene recombination checkpoint, and in mitochondrial function, among other processes – whose relationship to H3S10 biology is presently mysterious. **B**) H3S10 phosphorylation anticorrelates with transcription rate. TSS-aligned genes are aligned as in **Figure 3**, with TSS-aligned H3S10ph data being averaged for the indicated expression classes.

**Figure S3.**
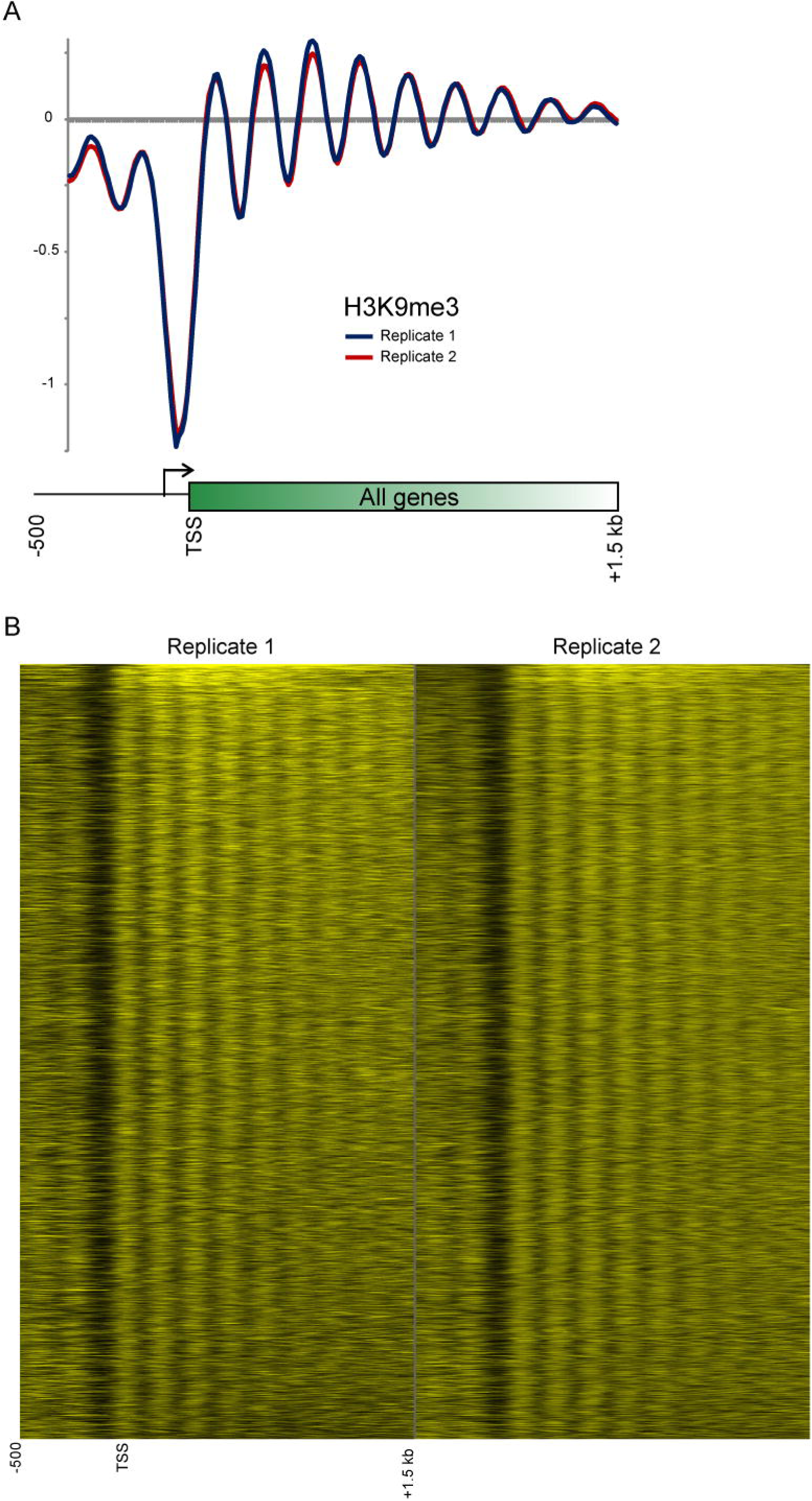
Replicate datasets for H3K9me3. **A**) TSS-aligned datasets for two independent biological replicates of H3K9me3 ChIP-Seq. **B**) Comparison between replicates of H3K9me3 ChIP-Seq, showing 2 kB surrounding the TSS for all ~5000 genes in *K. lactis*. For left and right panels, H3K9me3 signal is shown from black (zero reads) to yellow (saturated at 400 reads) color bar, for genes aligned by their TSS. Genes in both panels are in the same order, according to average H3K9me3 signal over protein coding region.

**Figure S4.**
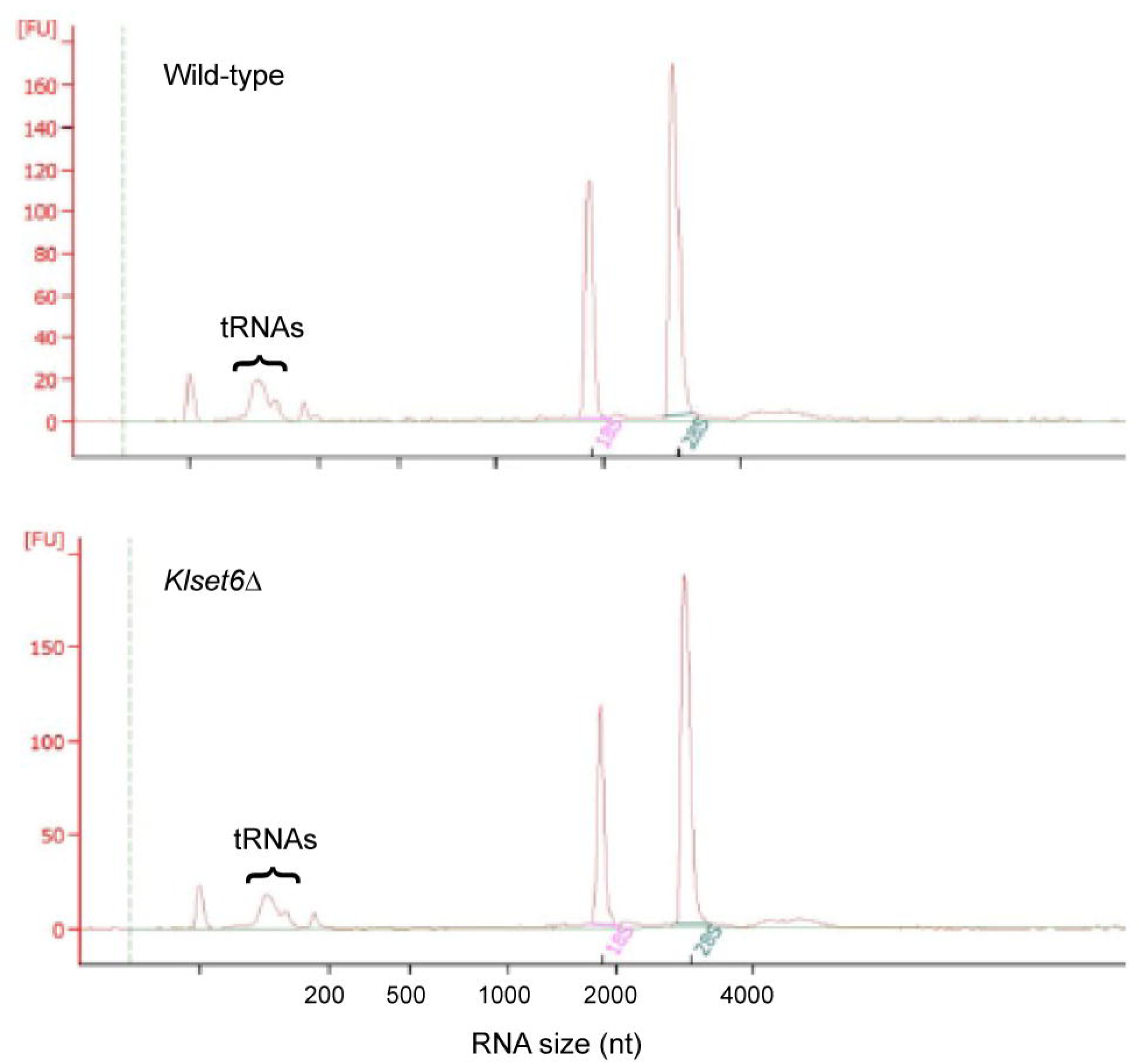
Total RNA profile from wild-type and *Klset6*Δ strains of *K. lactis*. Bioanalyzer RNA sizing for the indicated strains. Note that the fraction of total RNA in the 18S and 28S rRNA peaks, and in the tRNA size range, do not differ between the strains. Image is representative of two biological replicates, and separate analysis of small RNA (<300 nt) by Bioanalyzer confirms no bulk change in tRNA levels (compared to the 5S rRNA peak) in the deletion mutant (not shown).

